# Phoenix Enhancer: proteomics data mining using clustered spectra

**DOI:** 10.1101/846303

**Authors:** Mingze Bai, Chunyuan Qin, Kunxian Shu, Johannes Griss, Yasset Perez-Riverol, Weimin Zhu, Henning Hermjakob

## Abstract

**Motivation:** Spectrum clustering has been used to enhance proteomics data analysis: some originally unidentified spectra can potentially be identified and individual peptides can be evaluated to find potential mis-identifications by using clusters of identified spectra. The Phoenix Enhancer provides an infrastructure to analyze tandem mass spectra and the corresponding peptides in the context of previously identified public data. Based on PRIDE Cluster data and a newly developed pipeline, four functionalities are provided: i) evaluate the original peptide identifications in an individual dataset, to find low confidence peptide spectrum matches (PSMs) which could correspond to mis-identifications; ii) provide confidence scores for all originally identified PSMs, to help users evaluate their quality (complementary to getting a global false discovery rate); iii) identify potential new PSMs for originally unidentified spectra; and iv) provide a collection of browsing and visualization tools to analyze and export the results. In addition to the web based service, the code is open-source and easy to re-deploy on local computers using Docker containers.

**Availability:** The service of Phoenix Enhancer is available at http://enhancer.ncpsb.org. All source code is freely available in GitHub (https://github.com/phoenix-cluster/) and can be deployed in the Cloud and HPC architectures.

**Supplementary information:** Supplementary data are available online.

## 1 Introduction

Mass spectrometry (MS) has become the main technology in proteomics research and, consequently, proteomics data is growing rapidly. With more and more researchers sharing their data through public data repositories like PRIDE Archive (Perez-Riverol, et al., 2019) and iProX (Ma, et al., 2019), proteomics has become a “big data” field. As of 1 September 2019, PRIDE Archive hosts 13,389 datasets, representing 94,757 assays for acquiring proteomics data.

The number of unidentified spectra in public datasets (“dark matter”) is on average 75% of spectra measured in each MS experiment (Griss, et al., 2016; Perez-Riverol, et al., 2018). The main reason behind the low number of identified spectra, is that during the peptide identification step (Vaudel, et al., 2014) many derived peptides are either not present in the sequence database (e.g. sequence variants, or incomplete genome sequences) or they contain unexpected PTMs. In 2016, by clustering all PRIDE Archive data, we were able to identify almost 20% of previously unidentified spectra (Griss, et al., 2016).

In this manuscript, we presented the Phoenix Enhancer; a platform that enables researchers to perform comparison of their identified and unidentified spectra against previously published datasets in PRIDE Cluster. Four functionalities are provided: i) evaluate the original peptide identifications in an individual dataset, to find low confidence peptide spectrum matches (PSMs) which could correspond to mis-identifications; ii) provide confidence scores for originally identified PSMs, to help users to evaluate their quality (complementary to getting a global false discovery rate); iii) identify potential new PSMs for originally unidentified spectra; and iv) provide a collection of browsing and visualization tools to analyze and export the results.

### Design and Implementation

Phoenix Enhancer is designed for enhancing proteomics identifications by evaluating the quality of original PSMs or find new potential PSMs by searching the query spectra against a library of clustered spectra (Figure 1). It provides three core components, the front-end web interface, a Restful API and the data analysis pipeline. Users can upload the query files which include MS/MS spectra with/without identifications, and set the search parameters (**Supplementary information, Note 1)**. Then, the pipeline searches the query spectra against the spectral clusters and then score the previous PSMs and new recommend PSMs, finally write the scored PSMs into the MySQL database. The Restful-API or Phoenix Enhancer web can be used to retrieve or browse the final results. The analysis pipeline informs three major reports: i) potential incorrect identifications with a confidence score and a new suggested sequence if possible; ii) newly identified PSMs for previously unidentified spectra; iii) high confidence previous identifications which also got a high confidence score.

**Figure 1:**
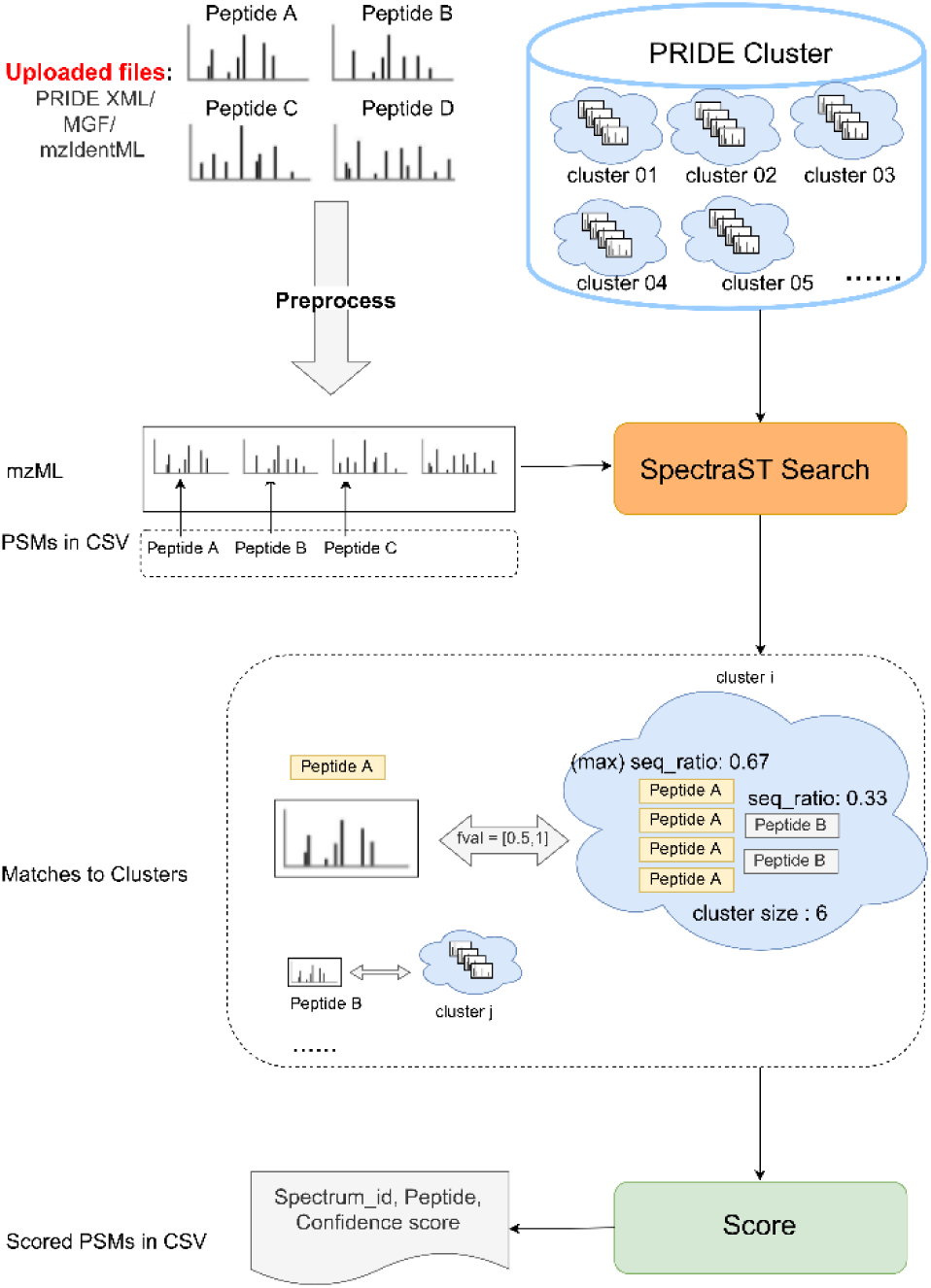
The Phoenix Enhancer Pipeline performs the actual analysis of the incoming spectra data by matching them to a spectral cluster archive and scoring the PSMs (original or recommend) by matched clusters.

#### Data analysis pipeline

When a file containing the MS/MS spectra with/without identificatons is uploaded, the Phoenix Enhancer pipeline converts the files to mzML; the MS/MS spectra are searched against the PRIDE cluster archives using SpectraST (Lam, et al., 2007) in “archive searching mode” (**Supplementary information, Note 2**). After picking the matches whose “fval” scores are higher than or equal to SpectraST’s default threshold (0.5), Phoenix Enhancer calculates the confidence scores for the 3 types of matched spectra/PSMs: previous PSMs get confidence scores for their previously assigned peptides; unidentified spectra or those in low confidence PSMs will get new recommend peptides with confidence scores, corresponding to the peptide sequence which has highest ratio within a cluster (called major peptide sequence).

Confidence scores can be used to assess the quality of the PSMs and to help the users finding novel PSMs. The detail of confidence score’s calculation is in **Supplementary information, Note 2.**

#### Restful API and Web

The interactive web interface and the restful API allow to: i) upload files for analysis, set analysis parameters; ii) show the results PSMs in tables and charts (**Supplementary information, Note 3**); iii) filter the results using species; iv) compare the query spectrum to matched cluster consensus spectrum (**Figure 2**); (v) check the details of the matched cluster, including comparing the spectra inside a cluster to its consensus spectrum; vi) download analysis result files for further analyses.

**Figure 2:**
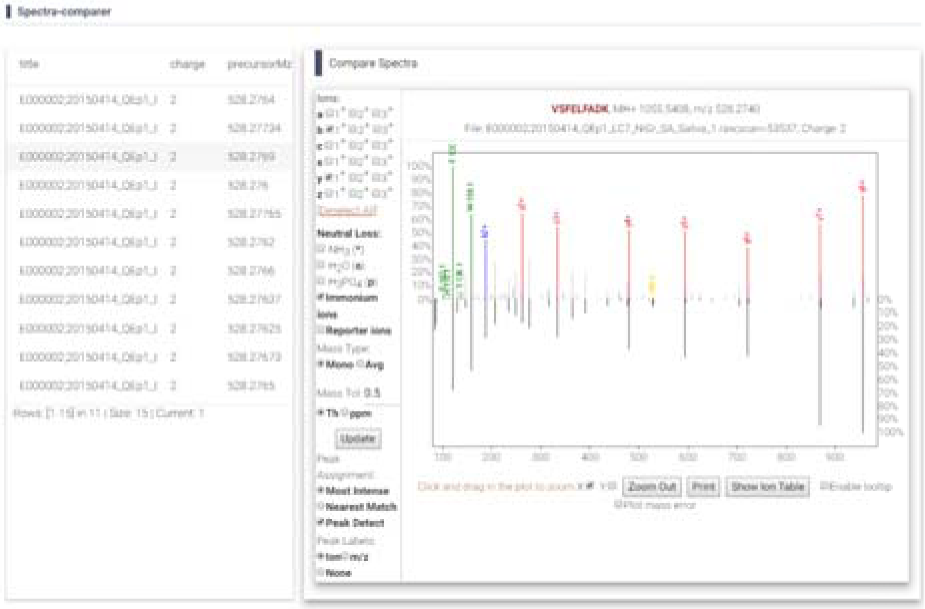
Spectra Comparer, for comparing query spectrum and matched cluster consensus spectrum. We have extended the original (https://github.com/UWPR/Lorikeet) spectrum viewer to provide the “butterfly” spectrum comparison facility.

In addition to the source code on GitHub, we provide our four components (web, web service, pipeline and MySQL server) as BioContainers images at Docker Hub (da Veiga Leprevost, et al., 2017).

### Benchmark datasets

We tested Phoenix Enhancer’s pipeline using 71 datasets from PRIDE Archive. 39 out of 71 (called “inside datasets” later) are already included in PRIDE Cluster, which are used to test the accuracy of the pipeline. We found that 0.172% of the previously identified spectra in all 39 “inside datasets” are incorrectly matched by the pipeline, and for 36 out of 39 “inside datasets”, the incorrect match rates are less than 1%. (**Supplementary information, Note 4**) Based on these figures, we believe the error matches given by SpectraST is in safe level and the results is acceptable

For 71 test datasets, we have found that the additional identifications range from 0 to 167%, the possibly incorrect original identification range from 0% to 5.27%, against the original number of PSMs. Interestingly, for datasets PXD000529, PXD000533, PXD000535; the original submissions contained 290 distinct phosphorylated peptides; while we can identify 1027 distinct new phosphorylated peptides. (**Supplementary information, Note 4**)

## Conclusion

In summary, we believe that Phoenix Enhancer is a valuable tool to take advantage of spectral clustering results for deeper proteomics data analysis, especially in validating the confidence of specific peptide biomarkers, and in finding interesting new potential biomarkers in repository datasets. The proposed Phoenix Enhancer is easy to deploy and reuse in local HPC and Cloud environments. The framework enables smaller research efforts to utilize the data mining potential of repository scale spectral clusters, and it provides new visualization, inspection, and validation tools for the results of such data mining efforts.

## Acknowledgments

This work was funded by the State Key Laboratory of Proteomics [SKLP-K201705] and National Key Research and Development Program of China (2017YFA0505002 and 2017YFC0906602].

