## Supplement Information for "Phoenix Enhancer: proteomics data mining using clustered spectra"

¬Application Note

**Motivation:** Spectrum clustering has been used to enhance proteomics data analysis: some originally unidentified spectra can potentially be identified and individual peptides can be evaluated to find potential mis-identifications by using clusters of identified spectra. The Phoenix Enhancer provides an infrastructure to analyze tandem mass spectra and the corresponding peptides in the context of previously identified public data. Based on PRIDE Cluster data and a newly developed pipeline, four functionalities are provided: i) evaluate the original peptide identifications in an individual dataset, to find low confidence peptide spectrum matches (PSMs) which could correspond to mis-identifications; ii) provide confidence scores for all originally identified PSMs, to help users evaluate their quality (complementary to getting a global false discovery rate); iii) identify potential new PSMs for originally unidentified spectra; and iv) provide a collection of browsing and visualization tools to analyze and export the results. In addition to the web-based service, the code is open-source and easy to re-deploy on local computers using Docker containers.

Menu

### 1 Start search on web

#### 1.1 Home page

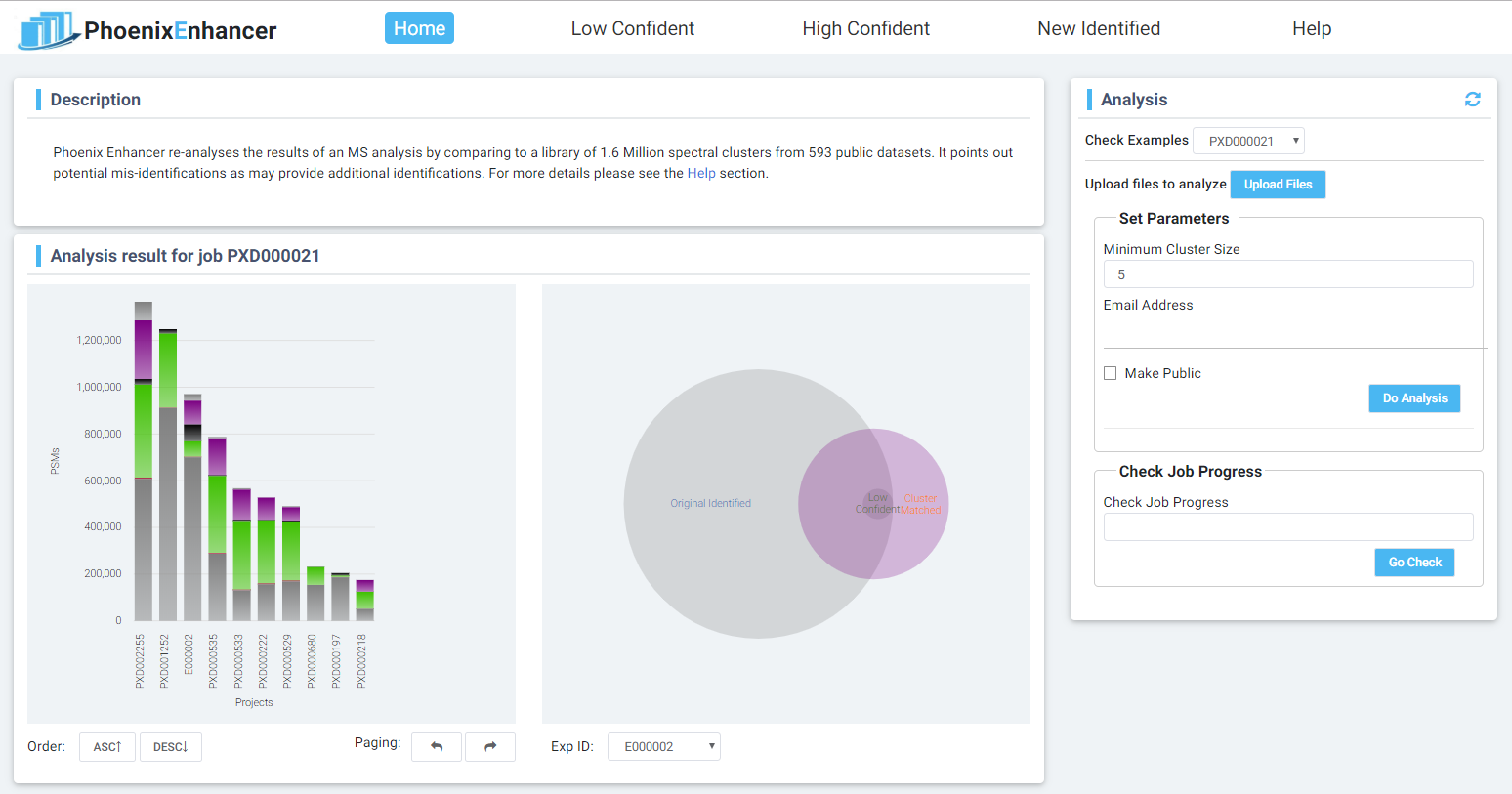

**Figure 1**. Phoenix Enhancer Home page.

Phoenix Enhancer is available at <http://enhancer.ncpsb.org>. At home page of Phoenix Enhancer, user is able to check public projects in the bar chart and Venn graph to see the statistics of those projects. Detail tables are accessed by clicking on the bar chart or the “Check Examples” selection. Besides, users can start their own analysis job at home page, in the right column. There are three steps to run an analysis job:

1. Upload MS/MS spectra data, which contains identification or not;

2. Set search parameters;

3. Start the analysis and check the progress.

Each of these steps will be demonstrated in below subsections.

#### 1.2 Upload files

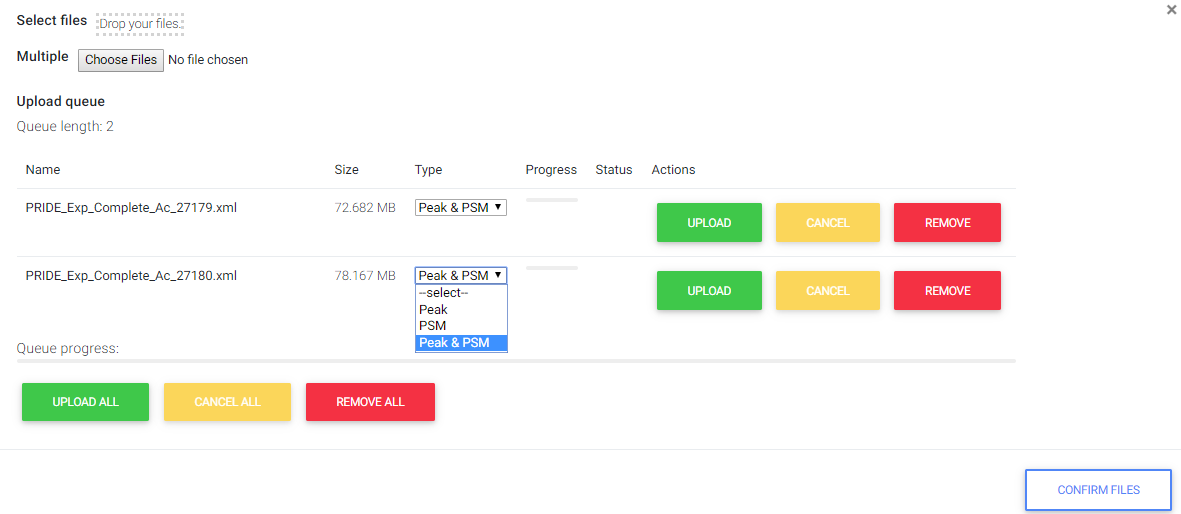

**Figure 2**. File uploading page.

Figure 2 shows the page of file uploading. Before the actually uploading starts, user should choose the type of files: “Peak” means only MS/MS spectra included, such as mzML, mzXML or mgf without SEQ section; “PSM” means only the matching information included, such as mzIdentML; “Peak & PSM” means both of them are included, such as PRIDE XML or mgf with SEQ section.

When the first file’s uploading starts, system will create an analysis job accession number and a random string token for this job. User will need them to access the result: private job is only accessible by token though public job is accessible for both accession number for and token.

After selecting and uploading files, user will still need to click the "CONFIRM FILES" button to confirm that all files have been uploaded and file types have been set. After confirming, user can’t delete, upload files or change the files’ type.

#### 1.3 Set search parameters

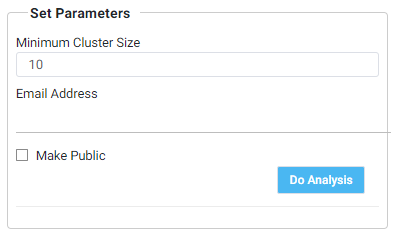

**Figure 3**. “Set Parameters” component.

Figure 3 shows the parameters for analysis job: minimum cluster size, email address of user, and to make this analysis job public or not. The “minimum cluster size” enables user to avoid matching to too small clusters which have low confidence to support the final confidence score. “User address” is needed to receive analysis report which contains job id, token and analysis result. If user agrees to make his/her analysis result public, he/she can tick on the "Make Public" button, to enable others see the analysis result at Phoenix Enahncer.

#### 1.4 Check pipeline status

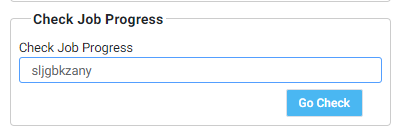

**Figure 4**. “Check Job Progress” component

Figure 4 shows the “Check Job Progress” component in home page. With the randomly generated token, user can check the progress in real time on the web page, by read the log file of the pipeline, as Figure 5 illustrated.

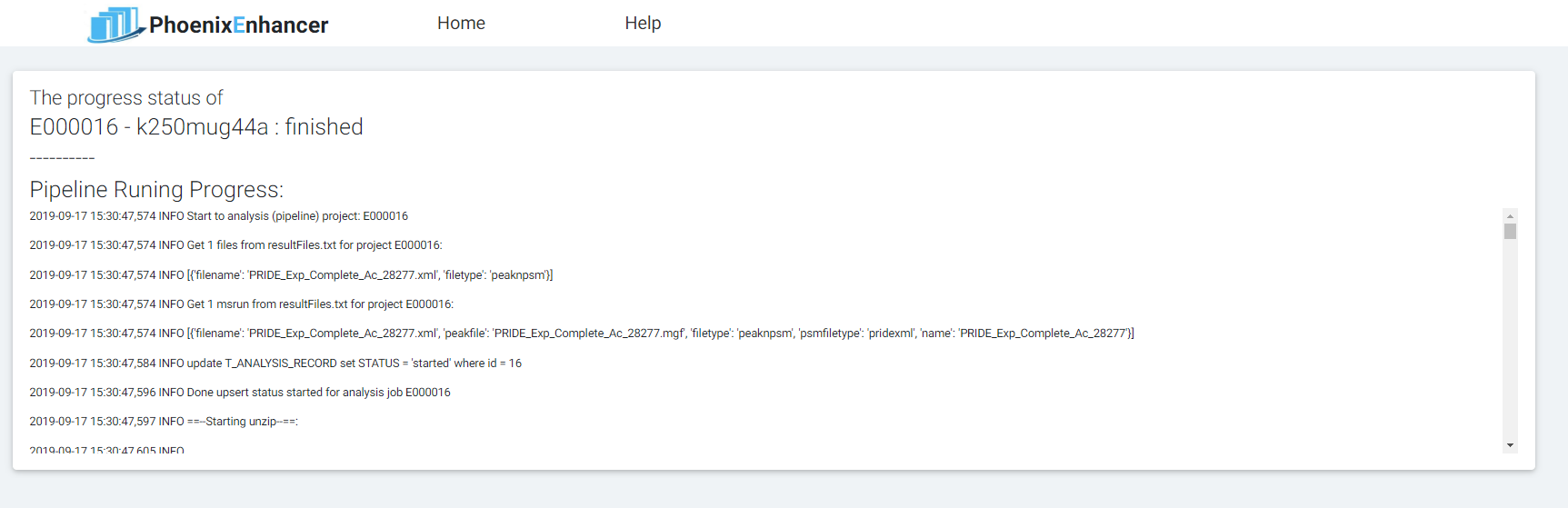

**Figure 5**. “Check Job Progress” page.

### 2 The pipeline

When a user is willing to evaluate the originally database search PSMs, one way is to cluster all these PSMs to all available identified spectra, or to the clusters in PRIDE Cluster[Griss, et al., 2016] or some other built spectra clusters, which is complicate and time consuming. Here we adopt another way, to address the clusters from PRIDE Cluster as spectral cluster archive and search the query spectra against it.

For these reasons below, we chose SpectraST for the pipeline to match query spectra to clusters: a). SpectraST provides options to enable users building the non-sequence spectral archive and to search the query spectra against it [Lam, et al., 2007]; b). The consensus spectra have been proved could match well in SpectraST, PRIDE Cluster’s consensus spectra were identified using SpectraST against the NIST human spectral library at a 1% peptide FDR in their test [Griss, et al., 2013; Griss, et al., 2016]; c). SpectraST is a widely used spectra library search engine which is integrated in TPP [Deutsch, et al., 2010]. SpectraST supports “spectral archive” building and searching from version 5.0 (released in Oct. 2016), which means the “spectral archive” be searched against only contains spectra (without the identified peptide information), suits our spectral cluster archive searching needs. SpectraST calculates dot product score between query spectrum and target spectrum in archive, then calculates a discriminant score *fval* based on the features such as dot product score of second hit and the number of matched peaks. In Phoenix Enhancer, a default *fval* cut off 0.5 is adopted.

Compared to SpectraST’s normal spectral library searching, Phoenix Enhancer’s methods have differences. In normal, the spectral library is built upon high confidence (the unique sequence in cluster) PSMs, spectra in library must be high-quality and be truly representative observations of the originating peptide ions in a relatively pure form, to ensure the accuracy and quality of the spectral library. For those highly similar spectra which are assigned completely different identifications by the sequence search engine, SpectraST will remove them all by multiple quality filters. In contrast, PRIDE Cluster keeps the original clustering information, which may help user to inspect the confidence level of this cluster, or other possible useful information such as the species distribution. With a filtering process similar to SpectraST, Phoenix Enhancer removes the little clusters which is smaller than 5 while SpectraST removes all the so-called “one-hit wonders” from the set of identifications to reduce the false discovery rates in large datasets. In addition, Phoenix Enhancer let users see and control the whole process of “spectral cluster archive” building.

After the spectral cluster archive searching, the confidence of PSMs (original and recommend) will be scored for sorting. While the confidence score is calculated based on these rules: i) each peptide in a cluster gets a confidence score, and the higher the peptide’s ratio, the higher the score; ii) for each peptide, if its ratio is higher than 0.5, it gets a positive score; if lower than 0.5, gets a negative score; iii) in addition to a peptide’s own ratio, the distribution of the other peptides’ ratios also matters: the more other sequences, or the more equally they are distributed, the higher the confidence for the major sequence; iii) the bigger the cluster size is, the higher the score, but we limit the weight of cluster size as 0.1 and use a cut off value at 1000 as the maximum cluster size. **Equation 1** is used to compute the score follow the rules above. The square root is used to measure how much the other peptides’ ratios affect the confidence of the considering peptide.

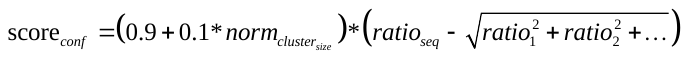
 **Equation 1**.

Then the scored PSMs will be write to MySQL tables for web browsing. The code of the pipeline is open resource and can be accessed at <https://www.github.com/phoenix-cluster/analysis-pipeline>.

### 3 Explore the results PSMs on web

#### 3.1 Three types of PSM tables

The mission of the web and web service is to provide an interface for user to start the analysis and to browse the analysis results: i) upload files for analysis, set analysis parameters and start the analysis (section 1); ii) browse the result PSMs in tables and charts; iii) filter the results using species; (iv) compare query spectrum to matched cluster consensus spectrum; (v) check the details of the matched cluster (will be represented in subsection 3.2); (v) export analysis result to files for further analyzing.

Figure 6 shows subtasks ii, iii, iv: Histogram charts shows the distribution of PSMs in confidence score (recommend confidence score for low confidence PSMs), cluster ratio and cluster size fields. Data table shows the PSM details (can be filtered by different species), user can mark the rows as accepted or rejected, based on the detail information. “Export to File” button for exporting and further analyzing (sub task v). The “Spectra-comparer” enable user compare the peak lists of query spectrum against the corresponded cluster’ consensus spectrum, accomplished with the matched ions (if available), etc.

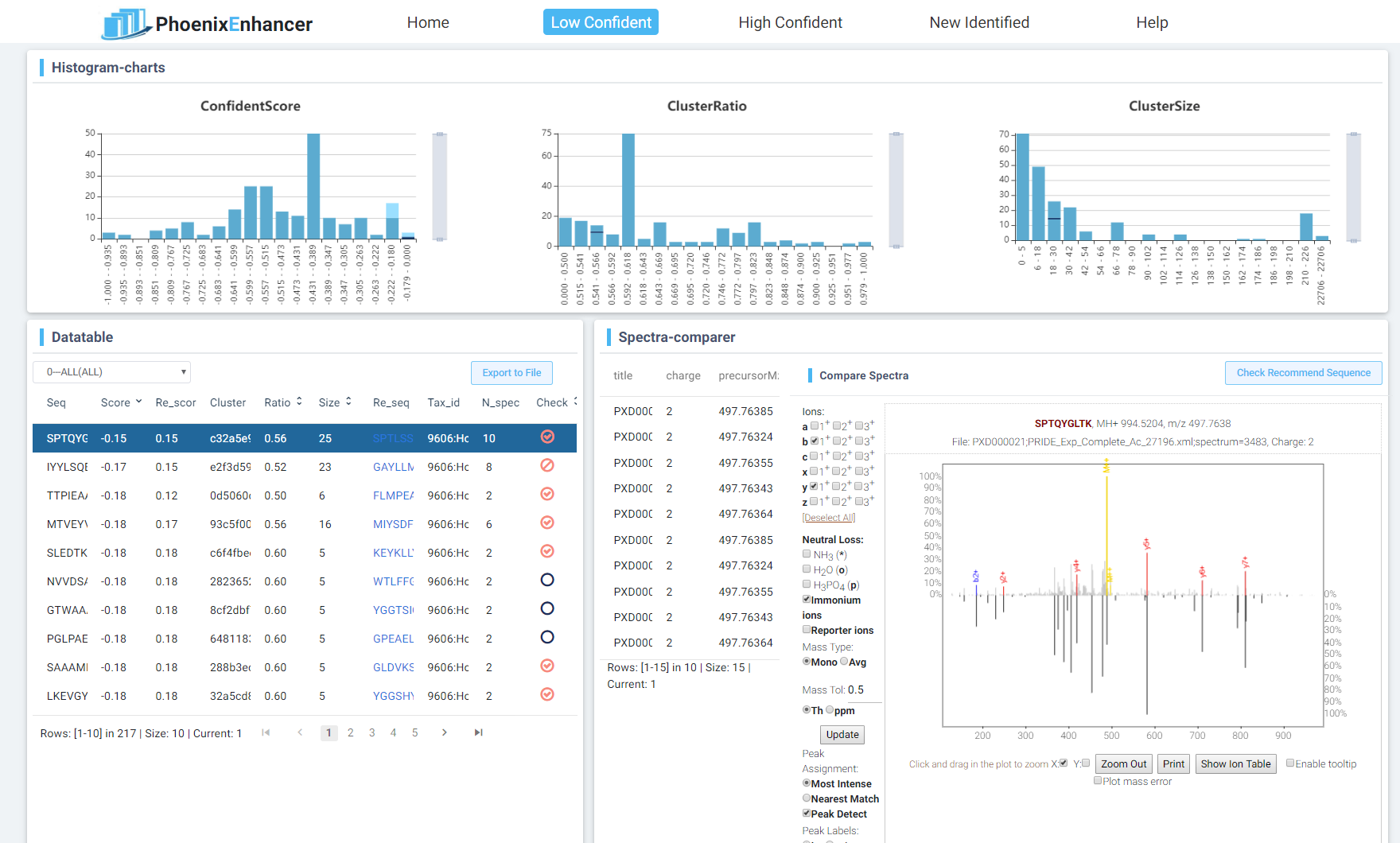

**Figure 6**. The result PSMs in tables and charts. The histogram charts show the PSM results’ distribution in confidence score (recommend confidence score for low confidence PSMs), cluster size and cluster ratio fields. Spectra-comparer, for comparing query spectrum to matched cluster consensus spectrum.

Angular 4 Framework (https://angular.io/) is the main framework used for implementing the web application. Spring Boot Framework (https://spring.io/projects/spring-boot/) is the main framework used for implementing the web service. The complete source code of the pipeline, web application and web service are available on GitHub (https://github.com/phoenix-cluster).

#### 3.2 Cluster details

As shown in Figure 7, by hovering on the cluster column in data table, user can get a pie chart showing the peptide sequence ratio distribution, for helping them to decide if the considering peptide sequence should be accept/reject with confidence. By clicking on the cluster id, web will jump to a cluster details page. Within this page (Figure 8), user reads more information about the cluster, such as the distribution of peptides, projects, as well as the clustered spectra within it. In addition, a spectra comparer is used here to compare the spectra in cluster and the cluster consensus spectra.

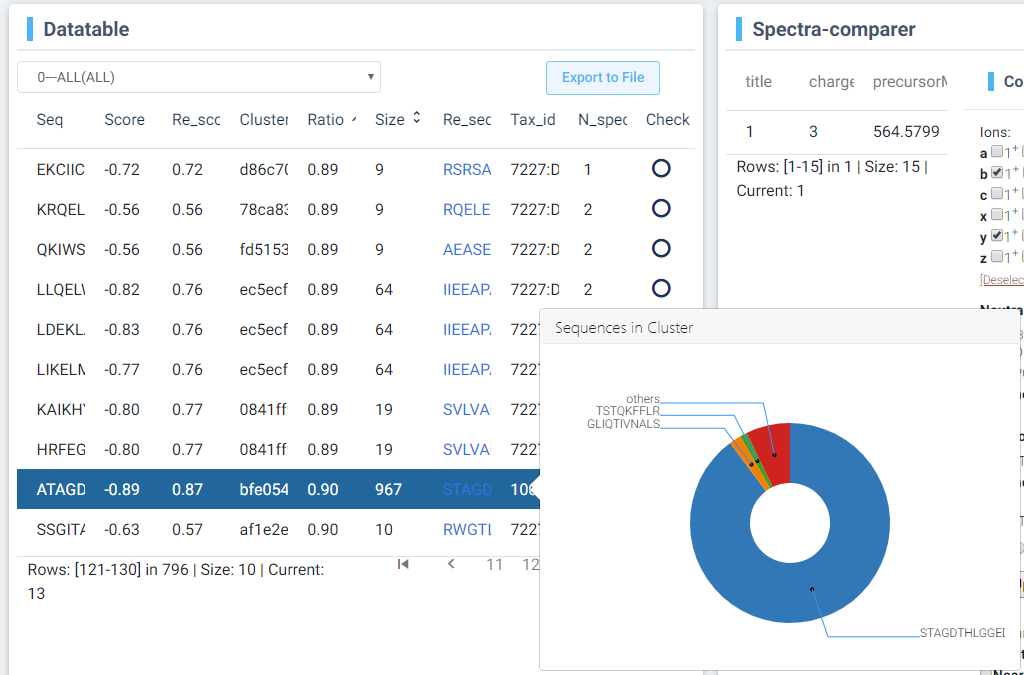

**Figure 7** The distribution of peptides in matched cluster, when hovering on the cluster Id.

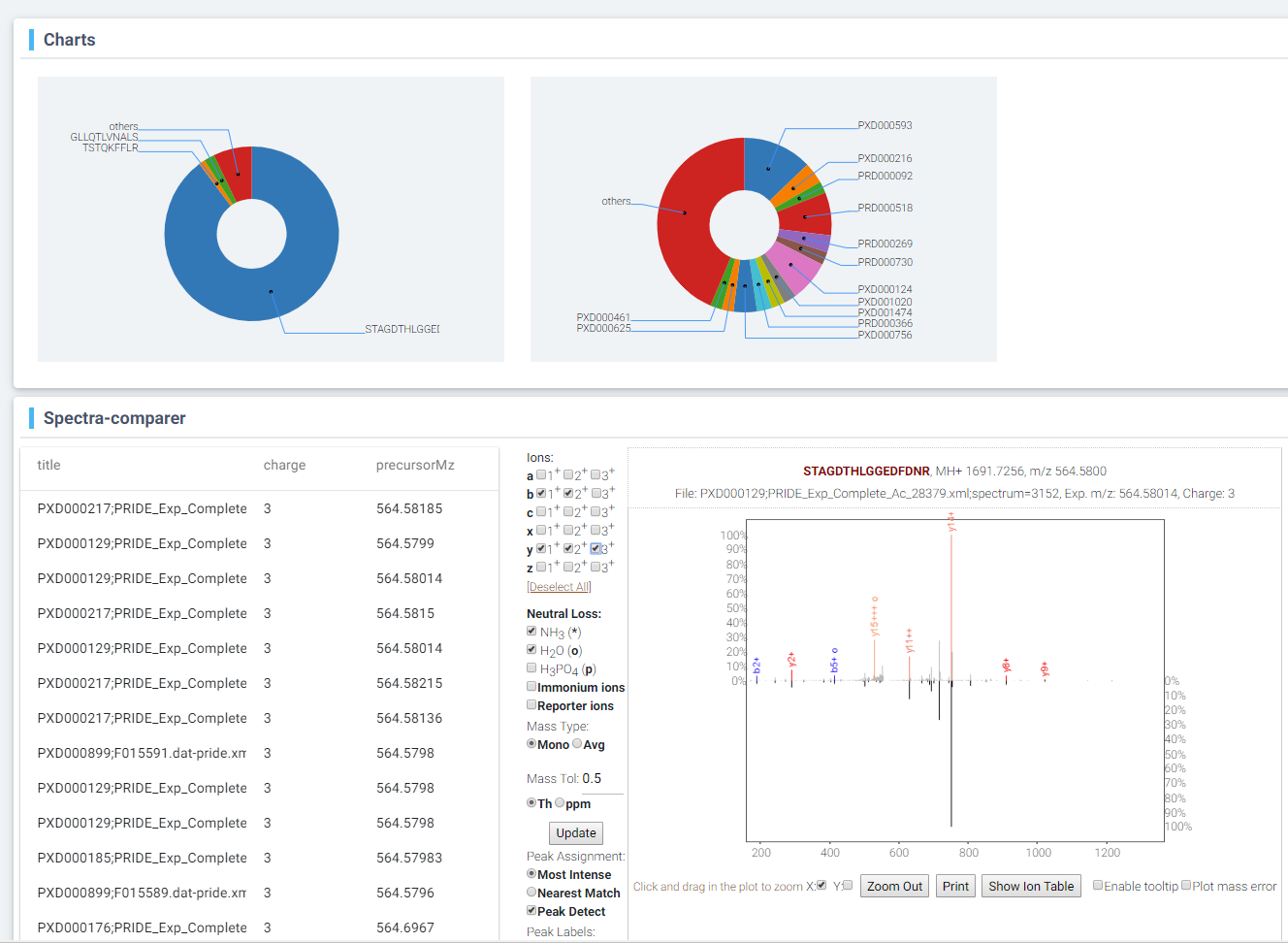

**Figure 8**. Cluster details page. The distributions of peptides, projects as well as spectra list are shown in this page, a comparer between the consensus spectrum and each of the clustered spectra is represented here too. Only the spectra stored in the database are available to show here. ([http:/enhancer.ncpst.org/cluster_details/bfe054f7-a371-4720-a673-f6f20dcb8e25](http://192.168.6.20:4201/cluster_details/bfe054f7-a371-4720-a673-f6f20dcb8e25))

### 4 Benchmark datasets

#### 4.1 Assess the accuracy of pipeline

For the 39 datasets which are already in PRIDE Cluster, called “inside datasets” later, we firstly collected spectra in these datasets, and matched them to the clusters in whole PRIDE Cluster by SpectraST (fval >= 0.5, cluster size >= 5). Then we calculated these figures: the number of spectra each of which matched to the original cluster it belongs (No.Original), the number of spectra matched to non-original clusters but with same sequence (No.SameSeq), the spectra matched to non-original cluster which has different sequence from its original cluster (No.DiffSeq), then we calculated the error rate as No.DiffSeq/(No.Original + No.SameSeq). As Table 1 shows, the error rate of most (36) “inside datasets” are less than 1%, and the error rate among the 39 inside datasets is 0.172%. Based on the figures, we believe the error matches given by SpectraST is in safe level and the results is acceptable.

**Table 1**. The accuracy assessment on the 39 “inside datasets”. 36 out of 39 datasets are less than 1% and the overall error rate is 0.172%.

| Project ID | No.Original/  No. Spectra Matched to Origin Cluster | No.SameSeq/  No. Spectra Matched to Cluster has Same Sequence | No.DiffSeq  No. Spectra Matched to Cluster has Different Sequence Cluster | Error Rate  No.DiffSeq/(No.Original + No.SameSeq) |
| --- | --- | --- | --- | --- |
| PXD000027 | 476 | 0 | 0 | 0.000% |
| PXD000053 | 4368 | 6 | 1 | 0.023% |
| PXD000070 | 22150 | 150 | 4 | 0.018% |
| PXD000095 | 843 | 2 | 0 | 0.000% |
| PXD000119 | 3 | 0 | 0 | 0.000% |
| PXD000129 | 56042 | 1836 | 358 | 0.619% |
| PXD000176 | 5774 | 220 | 45 | 0.751% |
| PXD000185 | 45740 | 469 | 104 | 0.225% |
| PXD000197 | 9065 | 142 | 567 | 6.158% |
| PXD000217 | 8506 | 541 | 102 | 1.127% |
| PXD000218 | 65723 | 844 | 84 | 0.126% |
| PXD000222 | 234222 | 1100 | 63 | 0.027% |
| PXD000312 | 16887 | 2834 | 39 | 0.198% |
| PXD000314 | 33267 | 210 | 1 | 0.003% |
| PXD000363 | 4152 | 515 | 10 | 0.214% |
| PXD000375 | 16603 | 38 | 9 | 0.054% |
| PXD000378 | 385 | 0 | 0 | 0.000% |
| PXD000384 | 32 | 0 | 0 | 0.000% |
| PXD000440 | 23 | 0 | 0 | 0.000% |
| PXD000450 | 5554 | 0 | 0 | 0.000% |
| PXD000464 | 409 | 9 | 9 | 2.153% |
| PXD000472 | 9160 | 59 | 0 | 0.000% |
| PXD000529 | 223934 | 2031 | 271 | 0.120% |
| PXD000533 | 262508 | 2930 | 130 | 0.049% |
| PXD000535 | 272746 | 5024 | 415 | 0.149% |
| PXD000558 | 11834 | 18 | 2 | 0.017% |
| PXD000674 | 2172 | 14 | 1 | 0.046% |
| PXD000688 | 707 | 4 | 0 | 0.000% |
| PXD000722 | 1366 | 8 | 0 | 0.000% |
| PXD000817 | 696 | 54 | 7 | 0.933% |
| PXD000878 | 12813 | 36 | 17 | 0.132% |
| PXD000899 | 7284 | 118 | 66 | 0.892% |
| PXD000948 | 32959 | 537 | 17 | 0.051% |
| PXD001099 | 4179 | 39 | 1 | 0.024% |
| PXD001118 | 23282 | 3400 | 126 | 0.472% |
| PXD001492 | 35456 | 300 | 46 | 0.129% |
| PXD001541 | 2152 | 318 | 6 | 0.243% |
| Total | 1433472 | 23806 | 2501 | 0.172% |

#### 4.2 Infer Identifications for Originally Unidentified Spectra

Previous works [Griss, et al., 2013; Griss, et al., 2016] have proved that an originally unidentified spectrum could be identified by clustering it into a reliable cluster, and our analysis in subsection 4.1 confirmed that our method can reliably assign the new coming spectra to clusters. This process is very similar with the spectral library searching process. Based on our test results on the 71 test datasets, the ratio of new identification number to the previous PSM number ranges from 0% to 167%. Details could be found at supplemental table 1 and figure 9. In figure 9, point color blue/gray means this project is/is not in the PRIDE Cluster. From this scatter line, we cannot see the existence in PRIDE Cluster affect the new identification rate obviously. From figure 9 we can also find that some projects can get high new identified PSM rate: 9 projects are higher than 50%, but the absolute numbers of previous PSMs are rather small. And for the dataset PXD001252, which has the highest number of previous identified spectra, no new PSM could be referred by Phoenix Enhancer at all. So far, we could find that the new identification rate is in a kind of randomly way, we can not predication how many new PSMs could be referred by our pipeline, however the average new identification rate is about 15.5%.

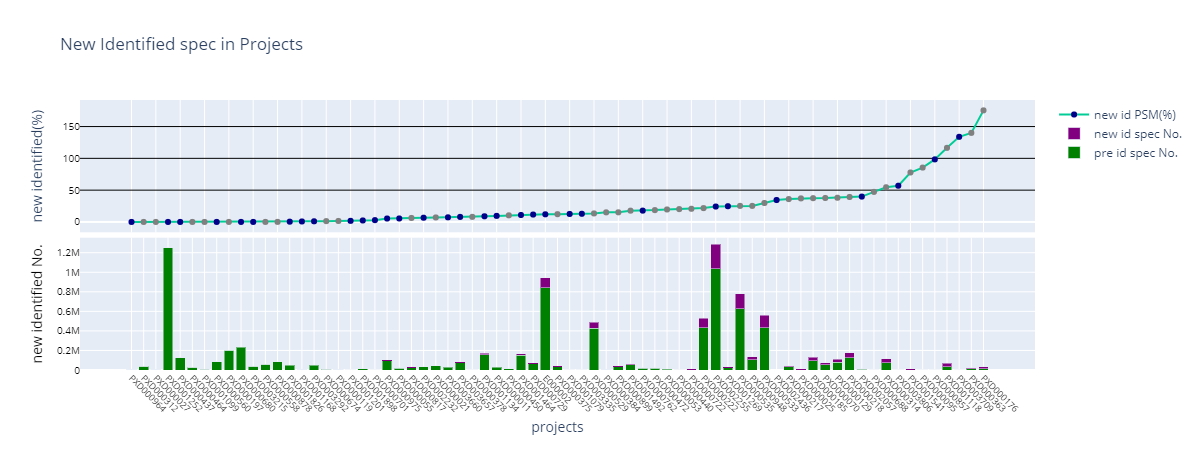

**Figure 9**. New Identified spectrum numbers in 71 test projects. Point color blue/gray means this project is/is not in the PRIDE Cluster. From the scatter line, we can not find the existing in PRIDE Cluster affect the new identification rate obviously.

For dataset PXD000529, PXD000533, PXD000535, the authors identified 290 distinct peptides with phosphorylation sites by clustering these datasets with phospho-enriched experiments, in publication [Griss, et al., 2016]. In our test, we can identify 1027 distinct new peptides with phosphorylation sites, has 222 distinct peptides overlap with their 305 distinct peptides. Some peptides (27) in the rest 83 peptides are in the PRIDE Cluster but not yet been matched here because they don’t have high similar spectra in the PRIDE Cluster.

#### 4.3 Identifying Common Incorrect Peptide Identifications/Better PSMs

Based on our test result on 71 datasets, the possibly incorrect original identifications be detected range from 0% to 5.27% and the average rate is 0.48%, against the number of original PSMs. For the datasets which are already in PRIDE Cluster, this rate is 0.78%; For the datasets which are not in PRIDE Cluster, this rate is 0.126%. The details are in supplementary table 1.

For dataset PXD003028 (as Phoenix Enhancer project E000002), we additionally identified 8089 peptide sequences (60050 spectra involved), which may come from 48 species. The results could be accessed at http://enhancer.ncpsb.org/new_id/E000002 or http://enhancer.ncpsb.org:8080/v1/scoredpsms/species/?identifier=E000002&Score%20type=posscore. Some of these species may not really in the sample, they have just been involved by non-unique peptides. Some of the 48 species have unique peptides in the additionally peptide set, such as Schizosaccharomyces pombe, which has not been included in the original sequence database. In the table of additionally identified peptides in Schizosaccharomyces pombe, we manually checked the results and got 17 unique peptides in Schizosaccharomyces (from 113 spectra) in PRIDE Cluster scope, then we searched these peptides on *unipept* (https://unipept.ugent.be/)[Gurdeep Singh et.al, 2018], 2 unique peptides were then found, see supplementary table 2.

To test if Schizosaccharomyces pombe is really in the sample, we reanalysis one sample’s data which has spectra matched to the cluster spectral archive as unique peptide in Schizosaccharomyces pombe (taxonomy id 4896). We used the parameters in Maxquant as same as the original searching to search against sequence database consists of UP000002485_284812.fasta and UP000005460_9606.fasta. We then identified about 20 proteins in the Yeast, with the FDR(QValue) threshold at 1%.

These results show that Schizosaccharomyces pombe or sibling species in same genus is possibly in this sample. In addition to the pipeline to match the spectra remained unidentified or are in low quality PSMs to detect originally undetected species, Phoenix Enhancer also provides the visualization tools to enable manual inspections and super links to *unipept* to check if these peptides are species unique.
